# A metagene of NRF2 expression is a prognostic biomarker in all stage colorectal cancer

**DOI:** 10.1101/690974

**Authors:** Séan M. O’Cathail, Chieh-Hsi Wu, Annabelle Lewis, Chris Holmes, Maria A Hawkins, Tim Maughan

## Abstract

**Objective:** Nrf2 overexpression confers poor prognosis in some cancers but its role in colorectal cancer (CRC) is unknown. Due to its role as a transcription factor we hypothesise a metagene of NRF2 regulated genes could act as a prognostic biomarker in CRC.

**Design:** Using known NRF2 regulated genes, we defined an NRF2 metagene to represent the pathway expression using principal component analysis and Cox proportional hazard models. The NRF2 metagene was validated in four independent datasets, including the recently profiled MRC FOCUS randomised controlled trial.

**Results:** 36 genes comprised the final prognostic metagene in the training set. 1,360 patients were included in the validation analyses. High NRF2 metagene expression is associated with worse disease free survival (DFS) and overall survival (OS) outcomes in stage I/II/III disease and worse OS in stage IV disease. In multivariate analyses, NRF2 expression remained significant when adjusted for known prognostic factors of adjuvant chemotherapy and stage in stage I/II/III disease, as well as BRAF V600E mutation and sidedness in stage IV disease. NRF2 metagene expression exhibits variation within each of the Consensus Molecular Subtypes (CMS) but high expression is particularly enriched in CMS 4.

**Conclusion:** We demonstrate in a large scale analysis that NRF2 expression is a novel biomarker of poor prognosis across all stages of colorectal cancer. Higher expression observed in CMS 4 further refines the molecular taxonomy of colorectal cancer.

## INTRODUCTION

Colorectal cancer is the 4^th^ most common cancer in the UK with 41,700 new cases per annum and only 57% of patients will live ten years or more [1]. An improved understanding of the biology of colorectal cancer may provide the basis for stratification of patients for differing treatment programmes, first by identifying prognostic effects. Current, known prognostic factors include ‘sidedness’ (left or right side of the colon) [2]—which is assumed to be surrogate of tumour [3] and patient biology [4]—RAS mutant status, BRAF status [5] and mismatch repair status [6,7]. The most widely used, RNA expression based classifier is the Consensus Molecular Subtype (CMS) [8]. It highlights that colorectal cancer is a significantly heterogeneous disease with different prognostic expectations among four subgroups. The taxonomy applied to each of the four CMS subtypes indicate enriched pathways, with no subtype defined by individual events, genetic aberrations or expression pathways.

The KEAP1-NRF2 pathway is a canonical stress response pathway which has shown prognostic importance in many tumours types, notably lung cancer [9]. NRF2 is a potent transcriptional activator that plays a central role in cell protection against oxidative and electrophilic stress. NRF2 activity is tightly regulated by KEAP1. Under basal (unstressed) conditions KEAP1, part of the Cul3 ubiquitin ligase family, mediates polyubiquitination of NRF2 protein [10]. During cellular stress changes in the thiol rich protein, KEAP1 modifies the structural integrity of the KEAP1-CUL3 ligase complex resulting in declining ubiquitination activity and an increase in cellular NRF2. NRF2 translocates to the nucleus and binds to antioxidant response element (ARE) sequences to regulate the transcription of suites of genes, including intracellular redox control, metabolic pathways, autophagy and drug transport [11]. Historically the NRF2 pathway was deemed to function in ‘tumour suppressor’ like capacity, allowing the cell to defend against stressors such as carcinogens [11]. However more recent evidence shows that some tumours acquire constitutive activation of the pathway which allow it to function in an ‘oncogene’ like fashion, promoting cell survival, resisting radiation, chemotherapeutics and dysregulating metabolism [12,13].

There are a number of distinct mechanisms by which the NRF2 pathway can become constitutively activated [14]. Mutations in both *KEAP1* and *NFE2L2* have been described in a up to 7.8% of colorectal cancers [15], although the rate in TCGA was less than 2.4% and 0.9% respectively [16]. Epigenetic modifications, methylation, of *KEAP*1 in colorectal tumours can silence its ability to regulate NRF2 [17]. Direct activation of NRF2 transcription via activated oncogenes such as *KRAS*^*G12D*^, *BRAF*^*V619E*^ and *c-MYC*^*ERT2*^ has also been described [18]. Indeed in CRC, the level of NRF2 signalling in the TCGA datasets is higher than would expect for the low somatic mutation rate observed [19], further suggesting complex post transcriptional mechanisms of activation.

Therefore it seems unlikely that *NFE2L2* expression in isolation will capture the full effect of pathway expression, and prove a useful biomarker. However, as NRF2 functions as a transcription factor controlling a battery of antioxidant response element (ARE) regulated genes, we hypothesise that a ‘metagene’ of NRF2 expression could be used to aggregate different mechanisms of pathway expression and act as a biomarker of prognosis in colorectal cancer. Here we detail for the first time the derivation of an NRF2 metagene from RNA expression data using a candidate gene approach [20] in colorectal cancer and demonstrate that high NRF2 expression is an independent biomarker of poor prognosis across all CRC disease stages.

## METHODS

### Candidate gene selection

Known NRF2 regulated genes were selected from two published prognostic lung cancer signatures [21,22] and refined for differential expression using the Oncomine database [23]. Input parameters “Cancer Type” and “Analysis Type” were set to ‘colorectal cancer and ‘cancer versus normal’ respectively. Differential expression was determined by threshold values of fold change > 2, p-value < 0.0001 and gene rank of top 10%. The database normalised gene expression across all selected datasets to allow summative gene expression comparisons. The resulting median gene rank for the meta-analysis across all selected datasets was calculated with its associated p-value, which was corrected for multiple hypothesis testing using the false discovery rate method [24].

### Construction of the NRF2 metagene

Principal component analysis (PCA) was applied to the probes representing the candidate NRF2 target genes. This generated a new set of continuous variables, principal components (PCs), which were weighted averages of the RNA expressions across the probes considered. Supervised variable selection was performed to decide how many major PCs^1^ would be useful for predicting prognosis in the training set. A Cox proportional hazard regression model was used to model the prognosis predicted by the PCs of NRF2 expression. The NRF2 metagene in each validation set was obtained by performing PCA on the corresponding probe sets (see supplementary figure 1 and supplementary information for further details).

### Datasets

Publically available colorectal datasets were downloaded from the Gene Expression Omnibus (GEO) database. Clinical and expression data was accessed from R programming environment using the packages ‘GEOquery’ [25] and ‘Biobase’ [26] obtained from https://bioconductor.org/biocLite.R. All datasets are summarised in Table 1. The datasets were divided into a training set, GSE17536 [27] and validation sets. Validation was carried out using datasets representative of non-metastatic, stage I-III disease (GSE14333 and GSE39582) [28,29], metastatic disease (MRC FOCUS trial) [30] and rectal only cancer (GSE87211) [31]. As part of the MRC Stratification in Colorectal cancer (S:CORT) consortium, we had access to the MRC FOCUS trial data including the RNA expression profiles generated by S:CORT (See supplementary information).

**Table 1.** Cohort characteristics of the training and validation sets. The numbers of cases, type of tissue, RNA expression platform, outcome variable and available covariates for adjusted analyses are indicated. (DFS = Disease Free Survival, OS = Overall survival, dMMR = deficient Mismatch repair, pMMR = proficient Mismatch Repair).

|  |  | **GSE17536**<br>**(Training**<br>**set)** | **GSE14333** | **GSE39582** | **MRC FOCUS**<br>**trial** | **GSE87211** |
| --- | --- | --- | --- | --- | --- | --- |
| **Patients** |  | 177 | 226 | 570 | 375<br>(355 DNA<br>mutant<br>status) | 189 |
| **Tissue type** |  | Fresh frozen | Fresh frozen | Fresh frozen | Formalin<br>fixed<br>paraffin<br>embedded | Fresh frozen |
| **Platform array** |  | Affymetrix<br>U133 v2.0 | Affymetrix<br>U133 v2.0 | Affymetrix<br>U133 v2.0 | Affymetrix<br>Xcel | Agilent<br>Human 4 x<br>44k v2 |
| **Primary site** |  | Colon | Colorectal | Colon | Colon | Rectum |
| **Stage** | I | 24 (13.6%) | 41 (18%) | 37 (6.5%) |  |  |
|  | II | 57 (32.2%) | 95 (42%) | 267 (47%) |  | 70 (30.8%) |
|  | III | 57 (32.2%) | 93 (41%) | 206 (36%) |  | 143 (63%) |
|  | IV | 39 (22%) |  | 60 (10.5%) | 375 (100%) | 14 (6.2%) |
| **Outcome variable** |  | OS | DFS | DFS<br>OS | OS | DFS<br>OS |
| **Covariates** |  |  |  |  |  |  |
| **Chemo(radio)therapy** |  |  | Yes 72 | Yes 240 | 375 (100%) | 189 (100%) |
|  |  |  | No 154 | No 326 |  |  |
| **Site of primary** |  |  |  | Prox. 232 | Left 203 |  |
|  |  |  |  | Dist. 351 | Right 152 |  |
| **BRAF V600E**<br>**mutation** |  |  |  | Mut 51 | 38 (10%) |  |
| **Mismatch Repair**<br>**status** |  |  |  | dMMR 77 | dMMR 15 |  |
|  |  |  |  | pMMR 459 | pMMR 326 |  |

### Statistical analysis

#### Primary analysis

The primary analysis for each validation set was to test whether the NRF2 metagene has a prognostic effect on disease free survival (DFS) and/or overall survival (OS). A Cox PH regression model with no variable was constructed, representing the null model. A second Cox PH regression model is fitted with the NRF2 metagene. The likelihood ratio test (LRT) is used to provide evidence against the null hypothesis (H0), that the NRF2 metagene provides no explanatory power. For the construction of Kaplan-Meier (K-M) curves, NRF2 metagene expression was subdivided by tertiles. Hazard ratios, and confidence intervals, presented for these curves are between the upper and lower tertiles. Significance testing was done only on Cox model analysis.

#### Secondary analyses

To assess whether the prognostic effect of the NRF2 metagene was confounded with other known prognostic variables we performed adjusted analyses using multivariate Cox PH regression models. Because the adjusting variables available varied across datasets, no adjusting variables were used in the training set. Prior to the adjusted analysis, the R package MICE [32] was used to impute missing values in the adjusted variables in GSE39582 and MRC FOCUS trial data. A LRT was performed to test the H0 that the model with only the adjusting variables adequately approximates the model with the adjusting variables and the NRF2 metagene. Significance suggests that NRF2 expression provides additional information to explain the survival outcome. The adjusting variables used for the secondary analyses are summarised in Table 1. All statistical analyses were conducted using the statistical computing software R [33].

## RESULTS

### NRF2 metagene derivation and training

In total 62 candidate genes were analysed in a total of 9 independent colorectal datasets for differential expression relative to normal tissue [16,34–39]. Some datasets were subsetted into different anatomical sites for the purposes of analysis resulting in 24 discrete sets of data (see supplementary Table 1). 40 were found to be differentially expressed, 21 of which were significantly over-expressed and 20 which were significantly under-expressed, in at least one or more of the datasets (Figure 1A). One gene, COL3A1, was shared as it was over expressed in some datasets and under expressed in others. Of the 40 differentially expressed genes, four could not be matched between the training and validation datasets so were omitted from further analysis. The final group of 36 genes was: *ABCA8*, *ABI3BP*, *ADAM12*, *ADRB1*, *ANGPT1*, *ANKRD29, ANKRD44, BCHE, C15orf48, COL3A1, COL5A1, EGLN3, LIFR, METTL7A, PCM1, PLAU, PLCB4, RECK, RGCC, RRM2, SEC14L4, SERPINH1, SFN, SLIT3, SPP1, TNS1, TOM1L2, TSPAN5, TTYH3, VSIG10, VCAN, AKR1C1, LRP8, NAMPT, PTGES, SLC27A5*. There was a very high level of co-ordinated expression between the majority of the 36 genes in the training dataset as evidenced by pairwise correlations (Figure 1B). This group of genes is designated the NRF2 metagene.

**Figure 1.**
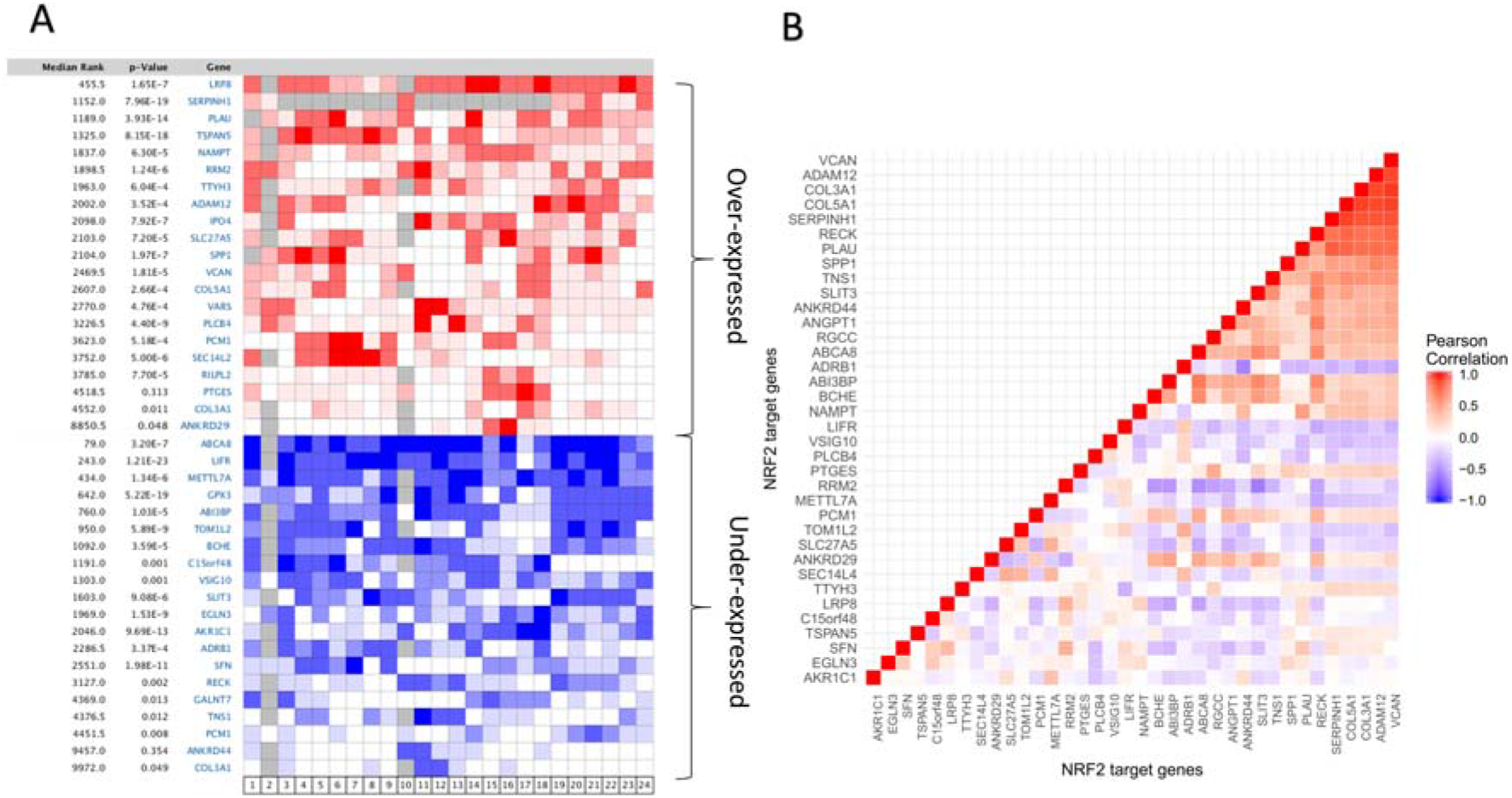
A) Aggregate heatmaps for Oncomine database over all over expressed and under expressed genes in 9 different colorectal datasets. The median rank for the meta-analysis and the corrected p-value are shown. 40 of the 62 candidate genes are significantly overexpressed and/or underexpressed in colorectal cancer relative to normal tissue at a FDR <0.05. 36 of these were represented in the GSE17536, after filtering by probe-matching, so were taken forward for further exploration. B) Pairwise correlation heatmap showing the degree of positive (red) and negative (blue) Pearson correlation between the 36 genes of the NRF2 pathway in the training set. The high positive correlation between a subgroup of genes (top right) indicates a high degree of co-expression.

#### Principal Component Analysis

In the training set the 36 genes are represented by 92 probes, where 65 of the 92 probes have an absolute correlation (in expression) >0.5 with another probe. This indicated potentially strong dependencies among the probes and so the expression levels measured by the 92 probes could be represented by a smaller number of variables. The 11 major PCs explained up to 80% of the variation in the 92 probes. The strong dependency among the probes was confirmed by the fact that the 81 minor PCs only explained <20% of the remaining variation.

#### Variable selection

AIC indicated that only the two major PCs, PC1 and PC2, were useful for predicting survival outcome, while BIC suggested that only the most variable PC (PC1) was useful. The inclusion of PC2 only slightly improved AIC, we therefore only considered PC1 in the training set. PC1 in the training set had absolute correlations >0.5 with probes that mapped to the following 10 genes: *VCAN, ADAM12, COL3A1, COL5A1, SERPINH1, RECK, PLAU, SPPI, TNS1* and *SLIT3*. Due to the high correlation of these genes with PC1 in the training set, we hypothesised that they were of higher biological relevance for prognosis prediction than other NRF2 target genes.

### NRF2 metagene a biomarker of worse survival

In stage I/II/III disease, higher NRF2 metagene expression corresponded to worse DFS in GSE14333 (HR^2^=1.551, 95% C.I 1.200–2.004; LRT, p = 0. 0008) and GSE39582 (HR=1.172, 95% CI 1.008–1.362; LRT, p = 0. 0383). Including the 60 cases of stage IV disease also available in GSE39582, NRF2 expression was also associated with worse OS (HR=1.240, 95% C.I 1.086–1.416; LRT, p = 0.001). In the MRC FOCUS trial, comprised of first line stage IV metastatic patients, NRF2 expression was again associated with a worse overall survival (HR=1.140, 95% C.I 1.035–1.255; LRT, p = 0.008). Figure 2 shows that high expression corresponded with worse prognosis for DFS in GSE14333 and GSE39582 (panels A and B), and for OS in GSE39582 and MRC FOCUS trial (panels C and D).

**Figure 2.**
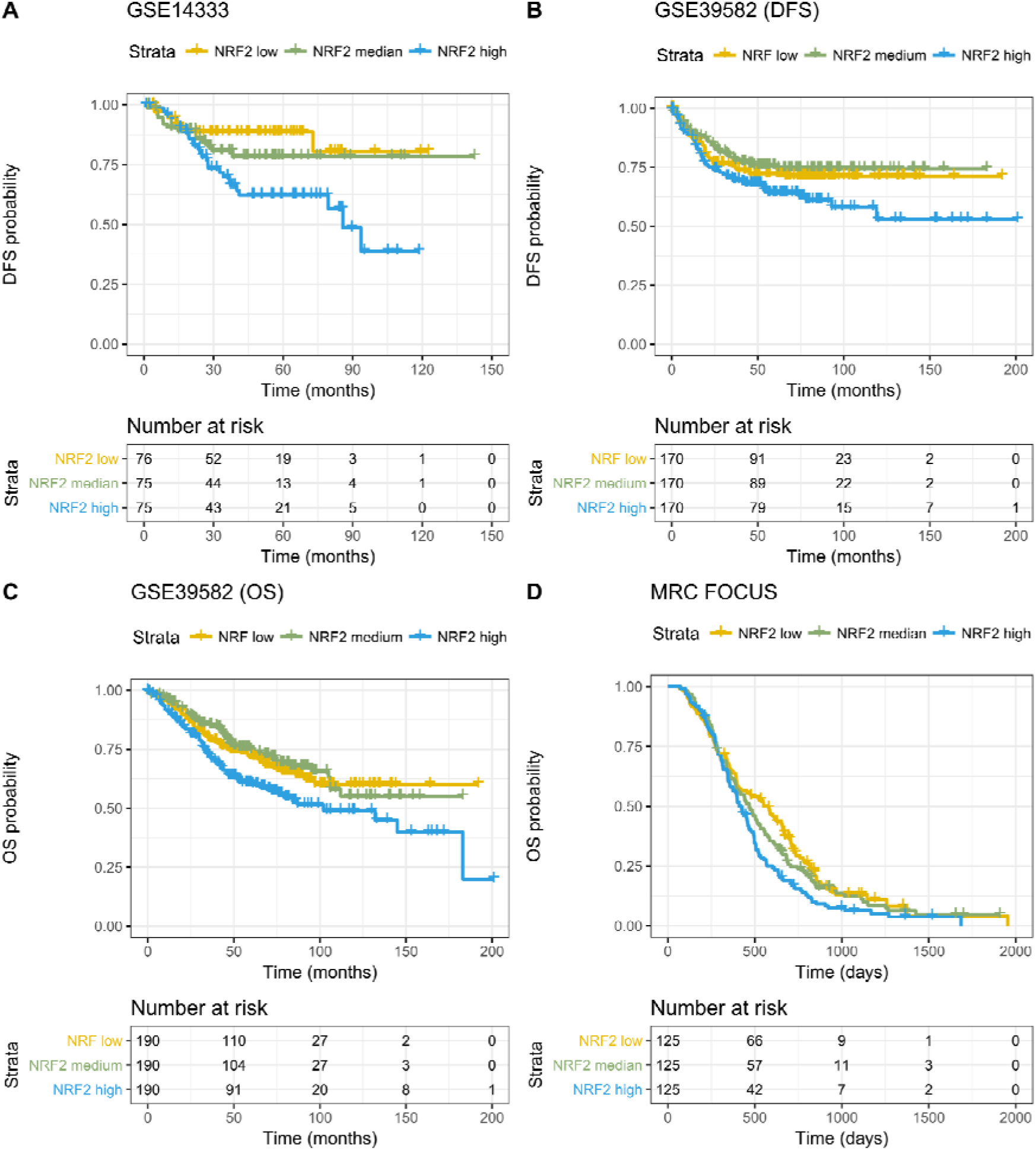
Kaplan-Meier curves and associated risk tables for the primary analysis of three datasets. All show survival outcomes for patients with high, intermediate and low (induced by tertiles) of NRF2 metagene expression. A) GSE14333 and B) GSE39582 represents early stage I-III patients C) GSE39582 represents stage I-IV patients and C) MRC FOCUS represents stage IV first line metastatic patients. For the Kaplan Meier curves, a median cut point was used to binarise NRF2 metagene expression.

In order to assess the relevance of NRF2 expression in rectal cancer specifically, and the ability to migrate between RNA expression platforms, we performed the analysis on a rectal cancer only expression dataset, where all sampled patients received neoadjuvant chemoradiotherapy (GSE87211). Higher expression was associated with worse DFS (HR=1.431, 95% C.I 1.060–1.933; LRT, p = 0.056) but not OS (HR=1.464, 95% C.I 0.955–2.245; LRT, p = 0.197). Figure 3 shows that high expression corresponded to worse prognosis for DFS.

**Figure 3.**
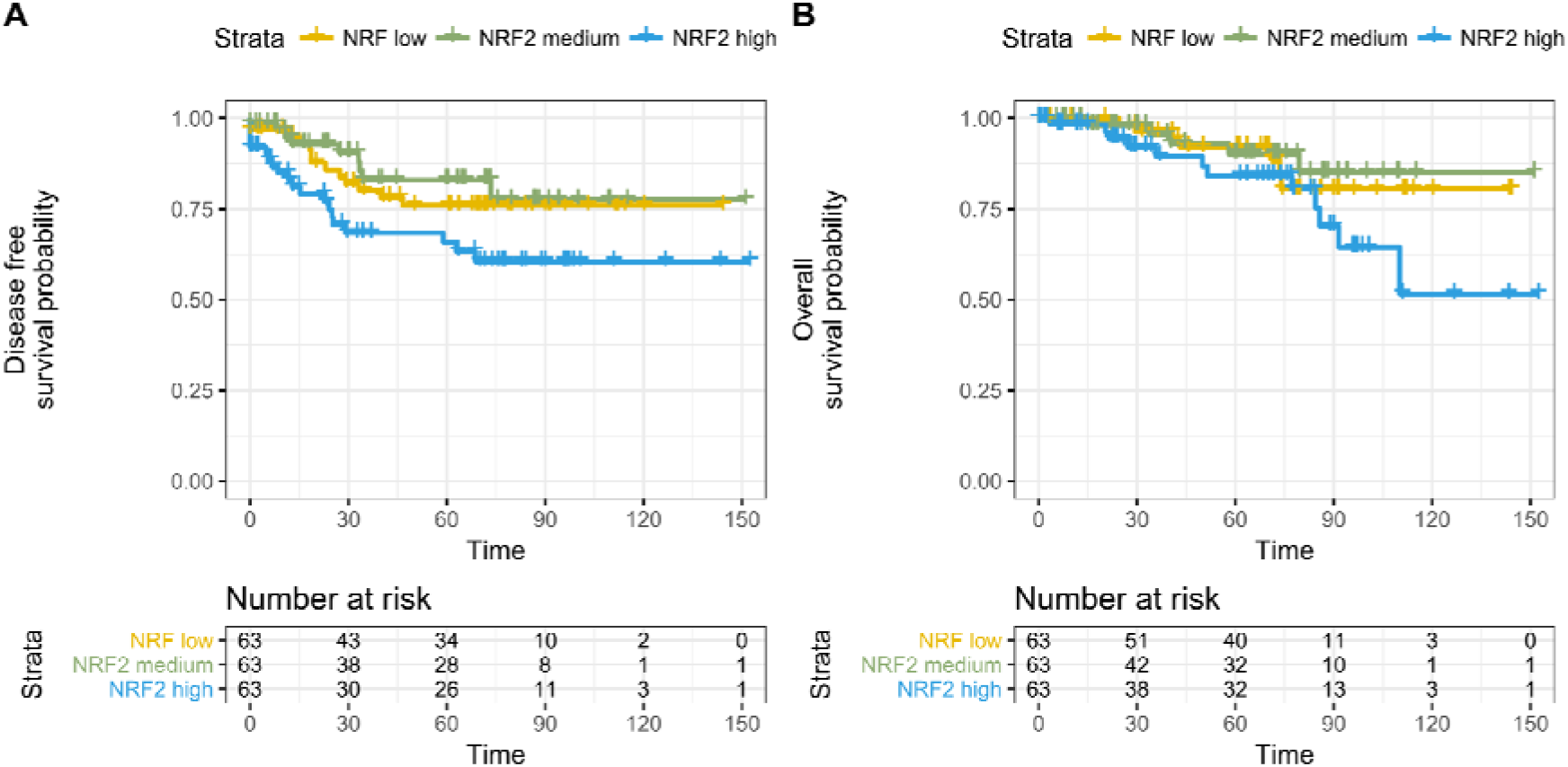
Kaplan Meier curves and associated risk tables for the primary analyses of GSE87211. On the left, high NRF2 metagene expression is associated with worse disease free survival with persistent separation of the curves. On the right, there was no effect on overall survival. For the Kaplan Meier curves, high, intermediate and low (tertiles) of NRF2 expression was used.

### NRF2 metagene provides additional explanatory power to known prognostic variables

Within the publically available datasets there were additional variables that are known prognostic factors. These magnitude of their respective effects are summarised in the forest plot (supplementary figure 2). We used these known prognostic factors in a multivariate analysis. In GSE14333, after adjusting for the effect of stage and adjuvant chemotherapy NRF2 expression remained a significant predictor of worse DFS (HR^3^=1.365, 95% C.I 1.049–1.776; LRT, p = 0.02). Similarly in GSE39582, the effect of high NRF2 expression corresponds to worse DFS (HR=1.168, 95% C.I 1.000–1.363; LRT, p = 0.049) after adjusting for the effect of stage and mismatch repair status (MMR). In the latter dataset, NRF2 expression was also significantly associated with worse OS when adjusting for stage alone (HR=1.185, 95% C.I 1.040–1.350; LRT, p = 0.01). No adjusted analysis was carried out for MMR with NRF2 expression on OS due to the known contrasting effects MMR status has on prognosis in early stage and metastatic disease, which could lead to model misspecification. In the MRC FOCUS trial, prognostic factors within the dataset were site of the primary tumour (sidedness) and *BRAF*^V600E^ mutation and used as adjusting variables in a multivariate analysis. Again, high NRF2 expression corresponds to worse overall survival (HR=1.123, 95% C.I 1.020–1.237; LRT, p = 0.0185). In summary, there was systematic evidence that NRF2 had an effect on DFS and/or OS in all available datasets (Table 2).

**Table 2.** Summary table of the Cox proportional hazard model analyses showing the numbers of patients included in each analysis, the adjusting variables where used and the p-value for the Cox model comparison (LRT = Likelihood ratio test)

|  | **N** | **Analysis** | **Variables** | **HR\***<br>**(C.I.)** | **LRT**<br>**(p-value)** |
| --- | --- | --- | --- | --- | --- |
| **Data set** |  |  |  |  |  |
| **GSE14333** | 226 | Primary | N/A | 1.113<br>(1.045–1.184) | 0.0008 |
|  |  | Secondary | Stage<br>Adjuvant<br>chemotherapy | 1.079<br>(1.012–1.150) | 0.0201 |
| **MRC FOCUS** | 375 | Primary | N/A | 1.033<br>(1.009–1.245) | 0.008 |
|  |  | Secondary | Sidedness<sup>‡</sup><br>BRAF<sup>V600E</sup><br>mutation<sup>‡</sup> | 1.029<br>(1.005–1.054) | 0.0185 |
| **GSE39582**<br>**(OS)** | 570 | Primary | N/A | 1.054<br>(1.02–1.089) | 0.001 |
|  |  | Secondary | Stage | 1.042<br>(1.010–1.076) | 0.01 |
| **GSE39582**<br>**(DFS)** | 510 | Primary | N/A | 1.04<br>(1.002–1.08) | 0.0383 |
|  |  | Secondary | Stage<br>Mismatch repair<br>(MMR)<sup>‡</sup> | 1.039<br>(1.000–1.080) | 0.049 |
| **GSE87211**<br>**(DFS)** | 189 | Primary | N/A | 1.127<br>(1.019–1.245) | 0.056 |
| **GSE87211**<br>**(OS)** | 189 | Primary | N/A | 1.135<br>(0.985–1.309) | 0.197 |

\* hazard ratio for event *per unit increase* in expression of NRF2 metagene (continuous variable) which is different to HR between the upper and lower tertiles
<sup>‡</sup> missing variables imputed

### NRF2 expression and Consensus Molecular Subtypes (CMS)

In order to understand how NRF2 expression aligns with the current transcriptomic landscape of colorectal cancer, we examined the distribution of the three groups of NRF2 metagene expression level across the four CMS subtypes in the MRC FOCUS trial (Figure 4). While high NRF2 expression can be seen across all subtypes, strikingly CMS 4 showed substantially higher NRF2 expression with no patients in the category of low NRF2 expression. By contrast, the majority of patients in CMS 2 or 3 had low and intermediate NRF2 metagene expression.

**Figure 4.**
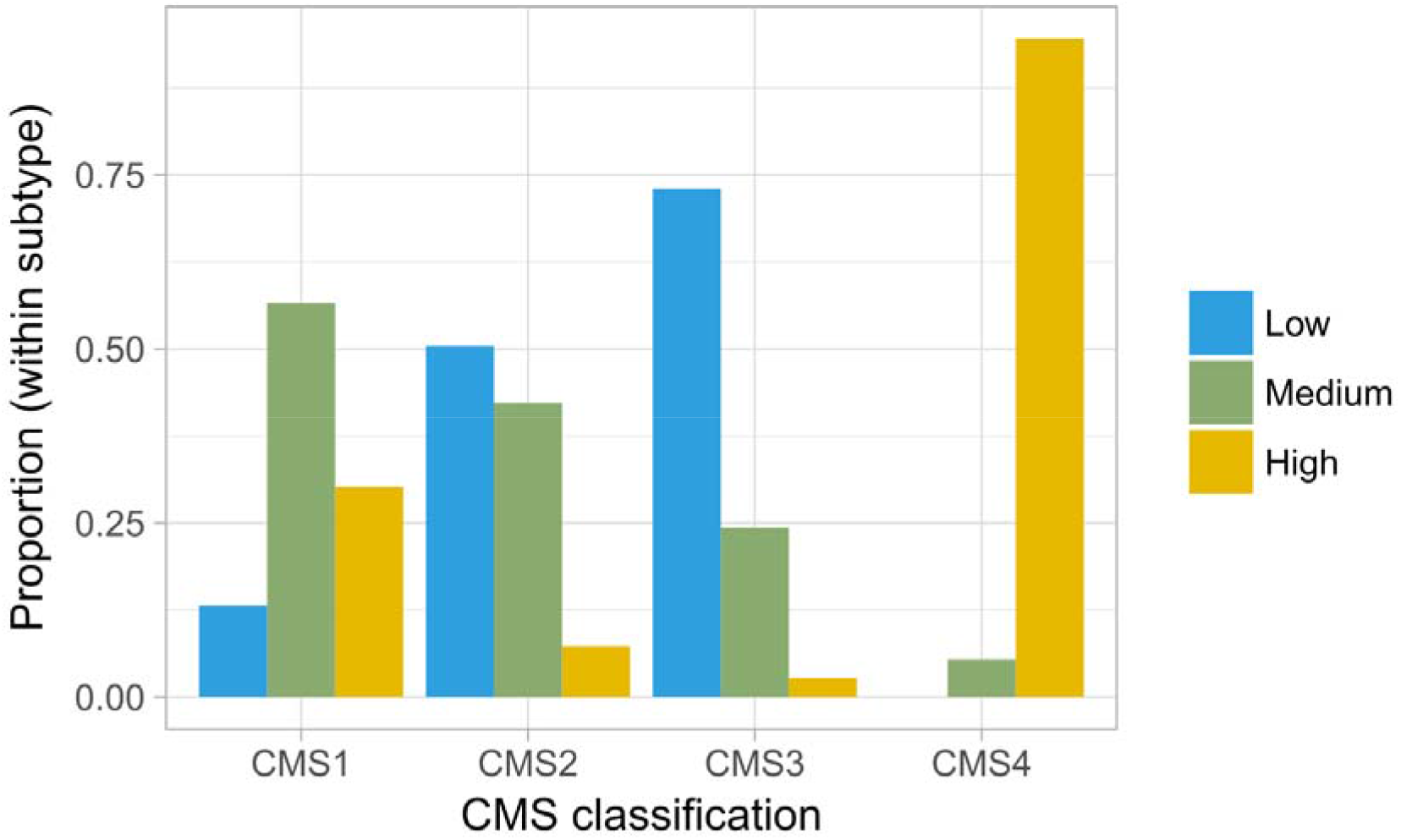
CMS classification was derived for the MRC FOCUS trial dataset. The barplot shows the proportion of high and low NRF2 metagene expression in each of the four CMS subtypes. CMS 4 was substantially enriched for high Nrf2 expression.

## DISCUSSION

In spite of the accumulating evidence that NRF2 plays a significant role in cancer [40,41] there is remarkably little data on the prognostic contribution of NRF2 in colorectal cancer. High levels of NRF2 expression within resected tumours have been found to be significantly correlated with p53 expression, Duke’s stage and poor clinical outcomes [42] as well as tumour size, TNM stage and metastases [43]. Whilst the latter investigated the association between NRF2 and survival status, their analysis did not take into account of the various aspects of time-to-event data, including data censoring or the concept of time. In addition, it was impossible to directly measure the prognostic effect of NRF2 in presence of potential confounders in their analysis framework. Therefore we set out to understand the effect of NRF2 expression in CRC and whether it provided additional information on prognosis to other established clinical and molecular variables.

We derived an NRF2 metagene to measure activation of the pathway in colorectal cancer and independently validated it as a biomarker of poor prognosis across all stages of colorectal cancer for the first time. This is an entirely new biological insight into colorectal cancer for several reasons.

To the authors’ knowledge, this is the first demonstration of NRF2 regulated genes behaving in a co-ordinated network fashion in CRC, *in vivo*, and supports our ‘metagene’ approach to represent KEAP1/NRF2 pathway expression. The high degree of co-expression seen between key genes *VCAN, ADAM12, SPP1, COL3A1*, *COL5A1, TNS1, SLIT3, RECK, PLAU* and *SERPINH1* was consistent with the predicted behaviour of these genes on the STRING database [44] (http://string-db.org/; supplementary figure 3). That the candidate genes were chosen on the basis of NRF2 regulation in lung cancer strengthens the unbiased nature of the analysis. With the exception of COL3A1, all of the key genes contain ARE within their promoter region emphasising NRF2’s ability to directly regulate transcription (supplementary information, supplementary figure 4).

Secondly, the NRF2 metagene has been shown as a biomarker of poor prognosis across all stages of colorectal cancer in several large, independent datasets comprising 1,360 patients making it one of the largest validation analyses of a transcriptomic biomarker in CRC to date. To place the analysis size in context, the OS and DFS cohorts used to validate CMS were 2,129 and 1,785 cases respectively [8].

Thirdly, the effect was detected from three different expression platforms using both FFPE and fresh frozen tissue. Lastly, and arguably most importantly, the prognostic effect is maintained when adjusting for known prognostic clinical and molecular factors including stage, adjuvant chemotherapy and mismatch repair status in the non-metastatic setting, and furthermore, *BRAF*^*V600E*^ mutation and tumour sidedness in the relapsed setting. The 375 patients with stage IV metastatic disease have been selected from a large randomised controlled phase III trial, which makes the findings highly stable and robust.

CMS has defined the current molecular taxonomy of CRC and is one of the most widely used RNA expression based biomarkers in CRC. The resulting classifications (CMS 1–4) and their respective prognostic outlooks describe the landscape in which any novel expression biomarker must be evaluated, especially as CMS was derived using a network-based clustering approach, agnostic of underlying biological mechanisms. We have shown that, although distributed across all four subtypes, high NRF2 expression is significantly enriched in CMS 4. This was unexpected. *A priori*, given that NRF2 was primarily known as a metabolic pathway and the literature supporting oncogenic KRAS activation of the NRF2 pathway, one could have expected enrichment within CMS 3, not CMS 4. High levels of NRF2 activation can promote angiogenesis and epithelial mesenchymal transition [41]. We would argue therefore that this adds further biological insight into the transcriptomic taxonomy of the CMS classification. The exact role of NRF2 in interacting with the current subtype annotations CMS 4 warrants further study.

The difference in prognosis between those with high and low NRF2 expression could, at least in part, be due to therapeutic resistance. Fluorouracil is the main chemotherapy drug used in both the adjuvant and metastatic setting, and was the backbone of therapy used in the FOCUS trial. Silencing of NRF2 signalling has been shown to overcome 5-FU resistance in colorectal cancer models in both an *in vitro* and *in vivo* setting [45]. Quantifying the effect of NRF2 expression in therapeutic resistance in relation to radiation and chemotherapy in colorectal cancer is ongoing in our laboratory. However, given the pluripotent nature of NRF2 it would seem plausible that it mediates poor prognosis by influencing multiple mechanisms [41].

Some limitations should be addressed. The amount of rectal tumours was low in the analysed data and the effect appears statistically less marked in the rectal only dataset GSE87211. However any argument that the finding is only pertinent to colon cancer ignores the fact that non-hypermutated colon cancers and rectal cancers are not distinguishable at the genomic level [16]. In fact, anatomical left colon more closely resembles rectum than right colon from a molecular biology standpoint. Furthermore, having accounted for sidedness and MSI where available, high NRF2 pathway expression remains a feature of poor prognosis. The statistical variance seen in GSE87211 is more likely the result of the relatively smaller number of events occurred (22% and 13% for DFS and OS respectively) which mitigated power. It may also due to an artefact of migrating between RNA expression platforms rather than a change in biological effect between colon and rectum.

Although we used a 36 gene ‘metagene’ to represent NRF2 pathway expression, NRF2 is known to regulate a large number of gene targets [46]. There may be an alternative group NRF2 targets which could better represent pathway expression in colorectal cancer. Fundamentally, the purpose of our analysis was representing NRF2 pathway expression from transcriptomic data to assess and understand its biological relevance in colorectal cancer. This is the first rigorous demonstration of its association with worse clinical outcomes in CRC.

The NRF2 metagene is a novel expression based prognostic biomarker, where higher expression associates with poorer prognosis across all stages of colorectal cancer. While the KEAP1-NRF2 pathway is increasingly well understood, better characterisation of its role and relationship to other biological factors in colorectal cancer is needed, particularly in relation to known CMS subtypes. The small number of genes needed to quantify metagene expression make it potentially suitable for development as a rapid diagnostic tool while its role as a canonical, cell intrinsic pathway makes it a relevant target for novel targeted therapies.

## Statements

### Contributions

Conceptualisation: SMO’C, CHW, AJL, MAH, TSM; methodology and data acquisition: SMO’C and CHW; data curation and statistical analysis: CHW; resources: SMO’C, CHW, AJL, CCH, MAH, TSM; writing (original draft): SMO’C, CHW; writing (review and editing): SMO’C, CHW, AJL, CCH, MAH, TSM; supervision: CCH and TSM.

## Supplementary Information

### Unsupervised dimension reduction

The metagene approach has the advantage of preserving statistical power. If all the candidate genes were included in an analysis, the statistical power would be exhausted. In addition, it was expected that some probes would be correlated due to multiple probes matching to the same gene symbol and co-expression of genes in the pathway. Such correlations create multi-collinearity in the downstream regression analysis, which leads to instability in parameter estimation. It was therefore desirable to obtain a simplified aggregate of NRF2 representation, which was achieved by applying principal component analysis (PCA) to the candidate NRF2 target genes.

PCA produced a new set of continuous variables, which are weighted averages of all the probe expressions. In the presence of high correlation among probes, the majority of the variation in the data can be explained by a smaller set of principal components (PCs). Thus PCA can provide a simpler representation of the expression data.

Specifically, we filtered out PCs that explained too little variation in the expression data and only retained the “major PCs”, which were the most variable PCs that explained up to 80% of the variation in the expression data. Such filter was applied because each of the excluded PCs would represent too little variation and would roughly be the same amount as that explained by other excluded PCs. Often, such minor PCs would have arbitrary probe weights, so should not be interpreted.

Furthermore, the PCs are uncorrelated among themselves, which eliminated the problem of multi-collinearity in the regression analysis downstream.

### Supervised selection of the principal components

Inclusion of major PCs that have little or no explanatory power can dilute the signal in an analysis. Therefore, supervised variable selection was performed to indicate how many major PCs were useful for and hence should be kept for prognosis prediction.

The Cox proportional hazard regression was used to model the prognosis prediction by the PCs of NRF2 expression. For variable selection, we progressively included PCs in decreasing order of variance. Akaike information criteria (AIC) and Bayesian information criteria (BIC) were calculated to indicate the most suitable number of major PCs to keep. The selected set of major PCs were used to represent the NRF2 metagene derived from the training set.

Prior to validation, we identified the genes that were represented by the most predictive PCs to understand the biological signal in the training set. If the absolute correlation of a PC and a gene’s probe expression was >0.5, then that PC would be considered to be representative of the gene of interest. For convenience, these genes are collectively referred to as the ‘represented gene set’. While a PC is a weighted average of the probe expressions of all the target genes that does not mean all the probes would be correlated with the PC. Therefore, the represented gene set would likely be a subset of the target genes.

In the training set, PC1 was the consensus between AIC and BIC. The represented gene set of PC1 were *VCAN, ADAM12, COL3A1, COL5A1, SERPINH1, RECK, PLAU, SPPI, TNS1* and *SLIT3*. These 10 genes were therefore deemed the best biological representation of the pathway that explained the survival outcome. Hence that was the pattern that should be sought following PCA in the validation sets, in order to ensure that we were validating the biological signal that was consistent with that discovered in the training set.

### Construction of the NRF2 metagene in the validation sets

The NRF2 metagene in each validation set was obtained by performing PCA on the corresponding probe sets. Major PCs were identified using the same filtering method for the training set. It is worth noting that the PCs are unsupervised summary statistics of the probe expressions. Thus, which PCs correlated with the probe expressions of a given gene could vary purely due to sampling variation. Specifically, while PC1 of the training set most strongly correlated with the represented gene set, a different set of PCs of a validation set might most strongly correlate with the represented gene set (derived from the training set). In that case, it would not be appropriate to use a fix set of PCs to validate the biological signal discovered in the training set.

For a given validation set, the correlations between the major PCs and the probe expressions of genes in the represented set were calculated. Based on the PC-probe correlations, PCs were selected to represent the metagene if those PCs were representative of any genes in the represented gene set. As discussed, the number of PCs selected via this approach could vary across datasets.

### RNA expression of MRC FOCUS

S:CORT undertook multiplexed analysis of a subset of FFPE primary tumour samples from the FOCUS trial (NCT00008060), which included RNA microarray analysis on the Affymetrix Xcel array. Samples were hybridised to the Affymetrix Xcel microarray as per the manufacturer’s instructions. Quality control analysis was run on samples using the R base ‘AffyQC’ module (https://github.com/BiGCAT-UM/affyQCModule). The files were then processed using the R packages ‘limma’ https://bioconductor.org.packages/release/bioc/html/limma.html) and ‘affy’ (https://bioconductor.org.packages/release/bioc/html/affy.html) to normalise the expression values using robust multiarray algorithm (RMA) and generate and expression matrix of probe intensities against the samples. CMS subtypes were determined from gene expression microarray data using the CMS classifier package in R (https://github.com/Sage-Bionetworks/CMSclassifier).

### Expression arrays and probe matching

To minimise differences between datasets induced by different platforms of RNA profiling, gene symbols were matched to probesets from the expression array annotation file, often resulting in multiple probes per gene. Probes were matched between platforms using genomic locus references from the annotation file (see supplementary table 2 – 4).

The RNA expression of GSE1433 and GSE39582 validation set was measured by the same platform, Affymetrix U133 v2, as that used in the training set so all probes could be used. However, the MRC FOCUS and GSE87211validation sets were respectively measured using the Affymetrix Xcel and Agilent Human 4 × 44K v2. In order to ensure that the metagene derived from the training can be translated to the IV validation set, we identified the U133 probes that matched to the Xcel and Agilent probes. Two types of matches were considered—(i) exact-match and (ii) sub/super-set-match. Exact match was where the chromosomal alignment from the annotation file of the microarray build probes matched perfectly. Sub/super-set-match occurred where the U133 probe was completely nested in an Xcel probe or vice versa. The same scheme was applied to matching to Agilent probes. In total the 36 genes were represented by 92 probes in Affymetrix U133 v2, 140 probes in Affymetrix Xcel array and 46 probes in Agilent Human 4 × 44K v2 array. Unmatched probes were filtered out prior to analysis. The 92 probe set in GSE17536 was used as the training dataset of this study.

### Regulatory sequence analysis

To further understand the role of NRF2 in regulating the represented gene set from the selected PCA in the training, *in silico* analysis of putative AREs in metabolic gene promoters were identified using RSAT [1]. The promoter sequence upstream to −5 kb lengths for named genes were retrieved from Eukaryotic Promoter Database (EPD) [2]. Subsequently, these sequences were analysed to identify AREs using the string-based pattern matching RSAT program ‘dna pattern’ [3]. The DNA patterns were entered as RTGASNNNGCR and RTGAYNNNGCR, where R = A or G, S = C or G, Y = C or T, and N = any nucleotide. These generic sequence queries were used in the query option to identify an exact match of an ARE sequence in the given 5-kb promoter sequences. The DNA patterns have been used previously [3] and are based on well-known previous publications on the consensus structure of AREs [4,5]. The analysis was carried out for *VCAN, ADAM12, COL3A1, COL5A1, SERPINH1, RECK, PLAU, SPPI, TNS1* and *SLIT3*.

---

**Supplementary Table 1.**
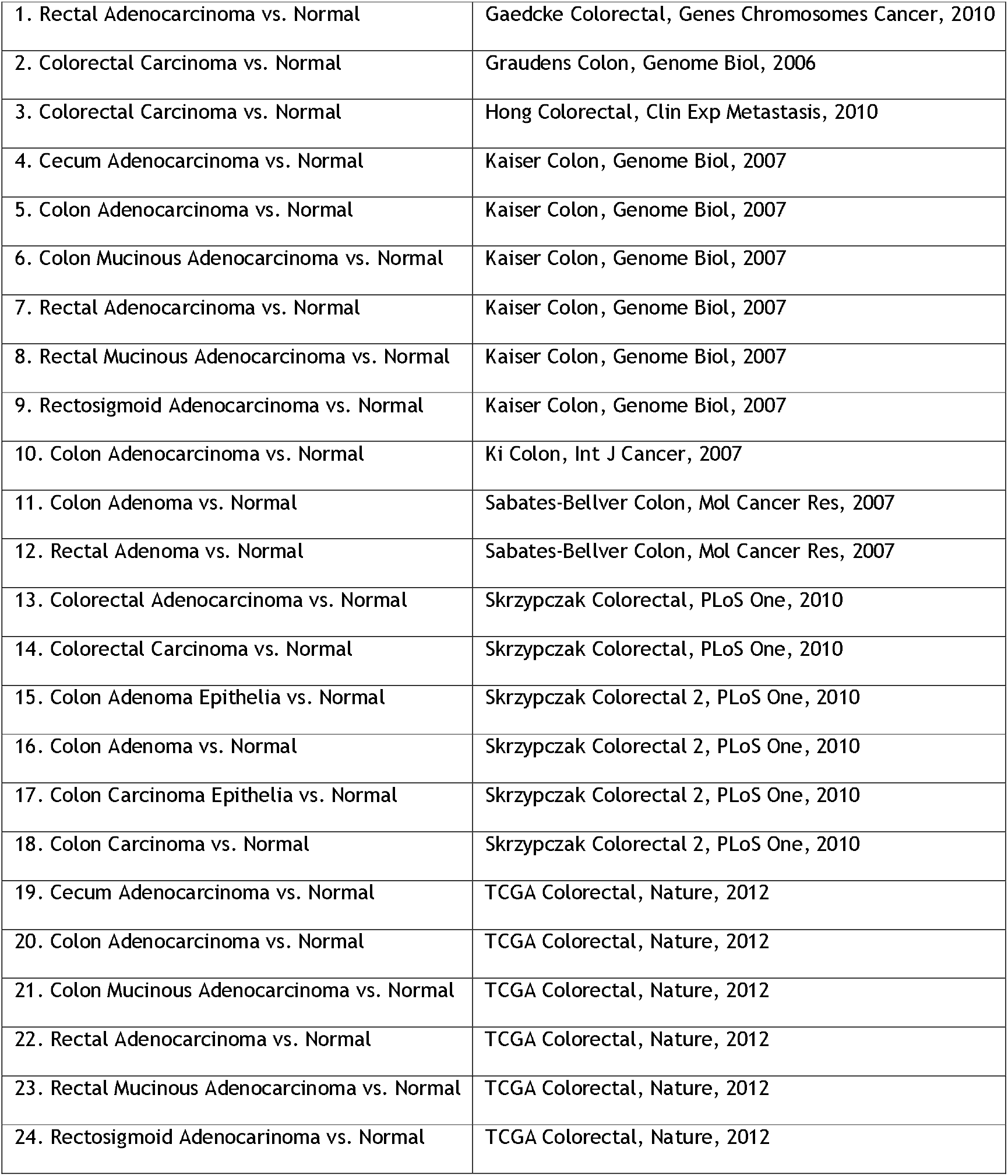
Description and source of the datasets selected in the Oncomine database to assess differential expression of the candidate genes

**Supplementary Figure 1.**
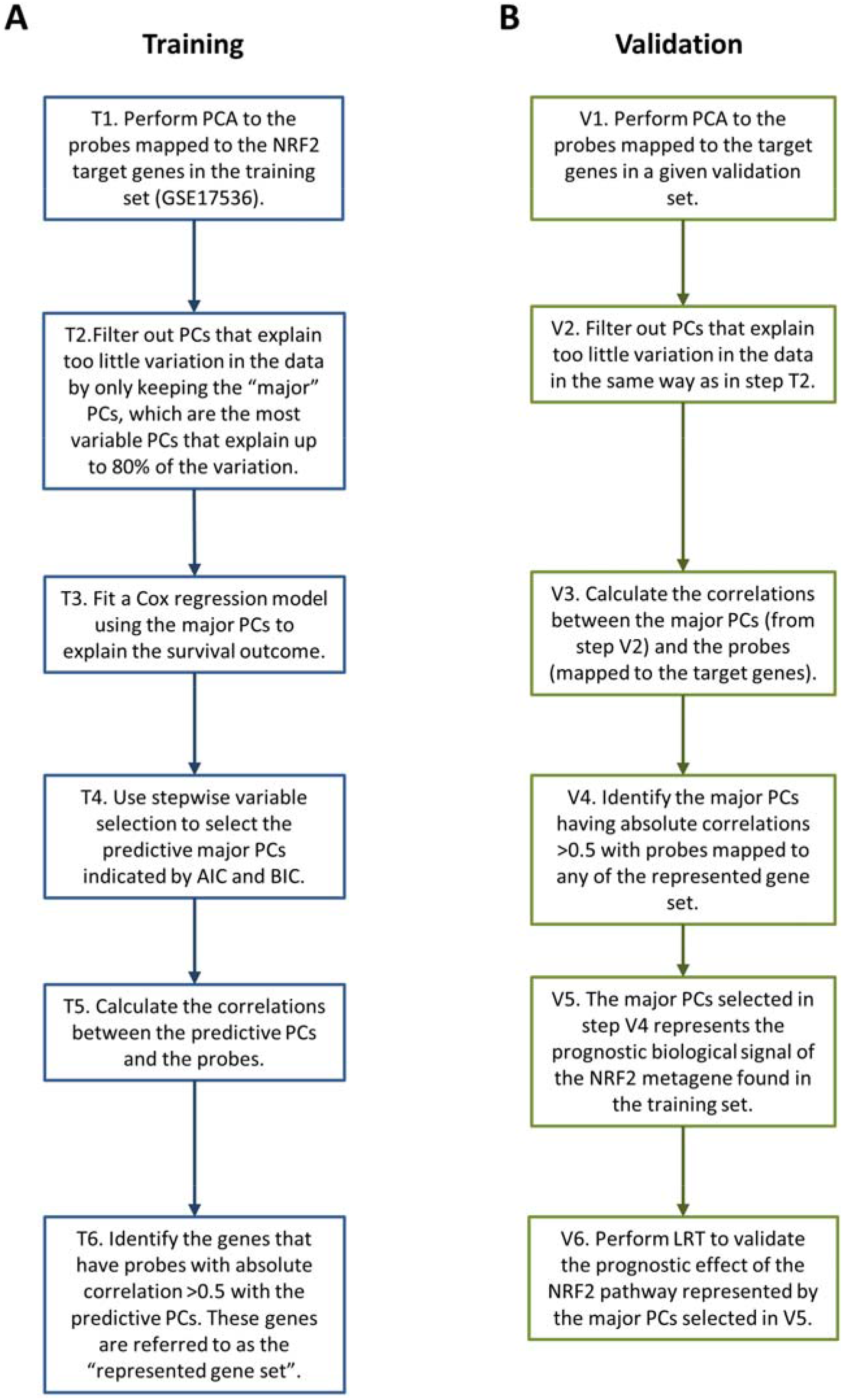
Flowchart describing A) training: the process of deriving the NRF2 metagene for prognosis prediction and B) validation: the process of constructing the NRF2 metagene for validation of its prognostic effect.

**Supplementary Figure 2.**
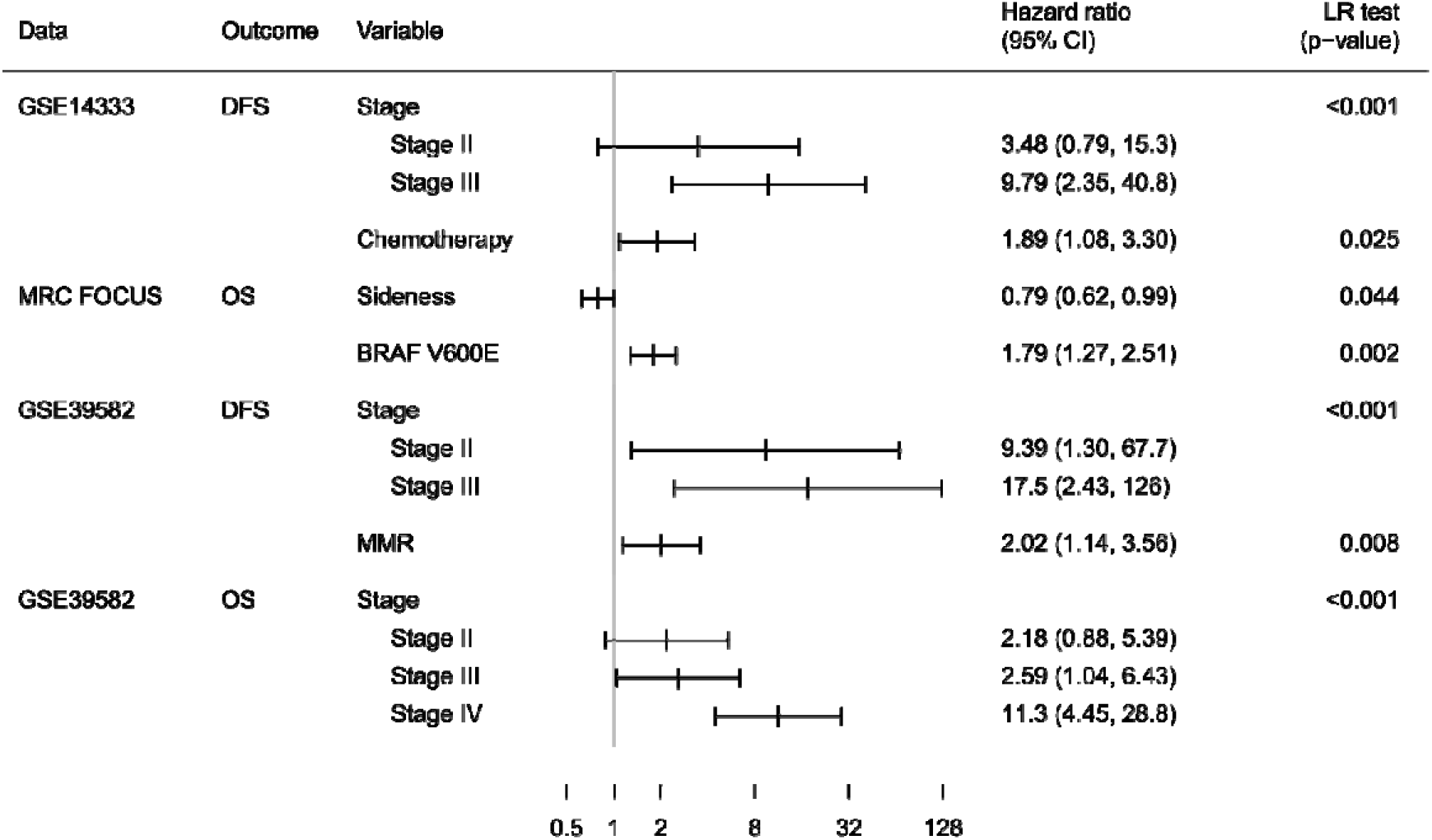
Forest plot showing the relative effects of the available prognostic variables within each dataset used in the validation analyses. These variables were used in the multivariate analysis with the NRF2 metagene.

**Supplementary Figure 3.**
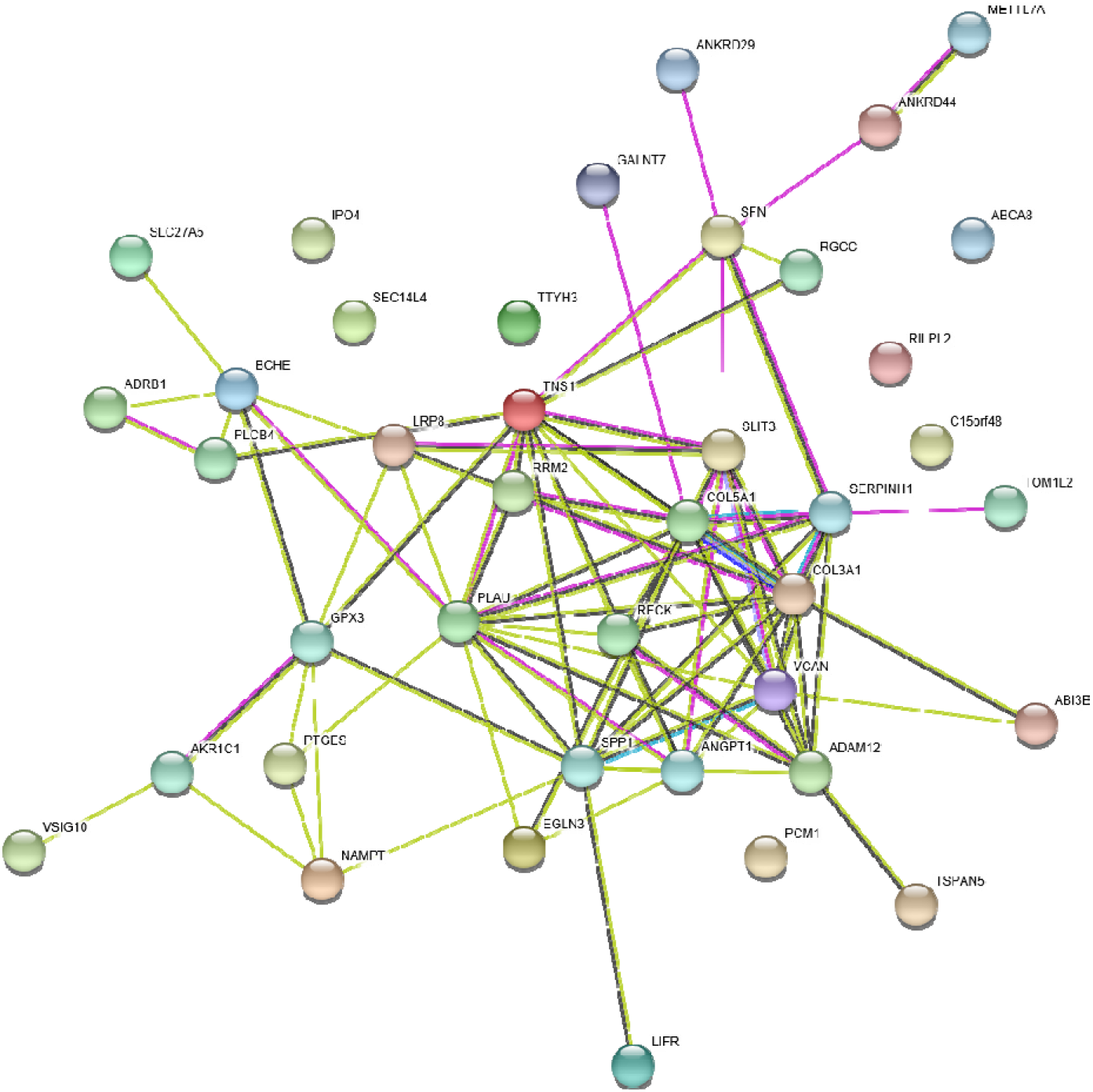
Known and predicted protein-protein interactions from the STRING database for the initial 40 genes from the refined differential expression analysis. The central network of genes (*VCAN, COL3A1, COL5A1, SERPINH1, RECK, PLAU, SPPI, TNS1 etc*) which show a high degree of connectivity are highly co-expressed in the training set. The line colour indicates the type of interaction and the thickness indicates the strength of interaction (thicker = stronger). [Cyan = known interaction from curated databases; purple = experimentally determined; green = gene neighbourhood; red = gene fusions; dark blue = gene co-occurrences; light green = textmining; black = co-expression; light blue = protein homology]

**Supplementary Figure 4.**
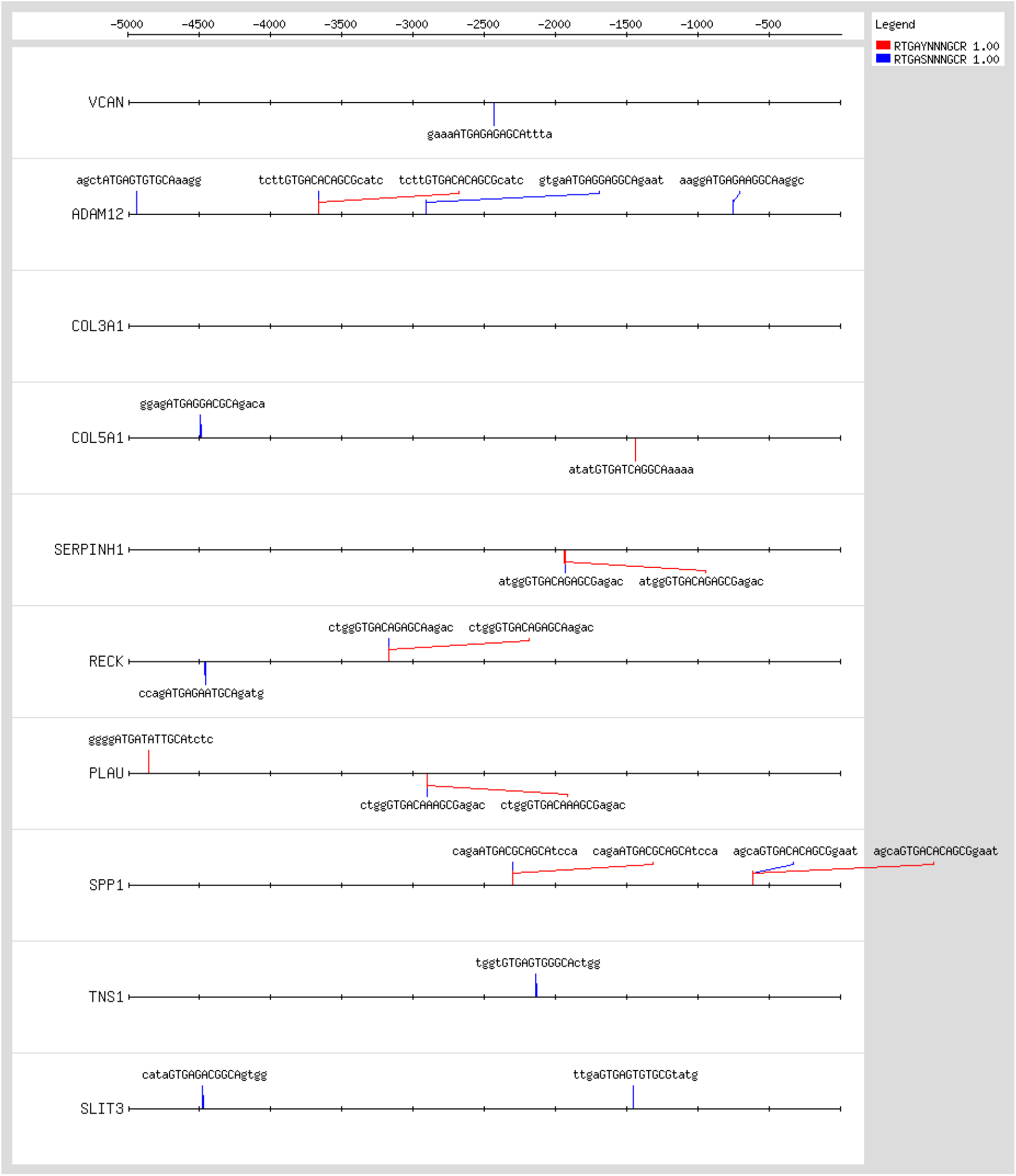
Regulatory sequence analysis of the 5000 bp region upstream of the start site of the representative 10 genes which have a high correlation with PC1 in the training set. 9 of these genes contain consensus ARE sites for NRF2 to bind and directly regulate gene transcription.

**Supplementary Table 2.**
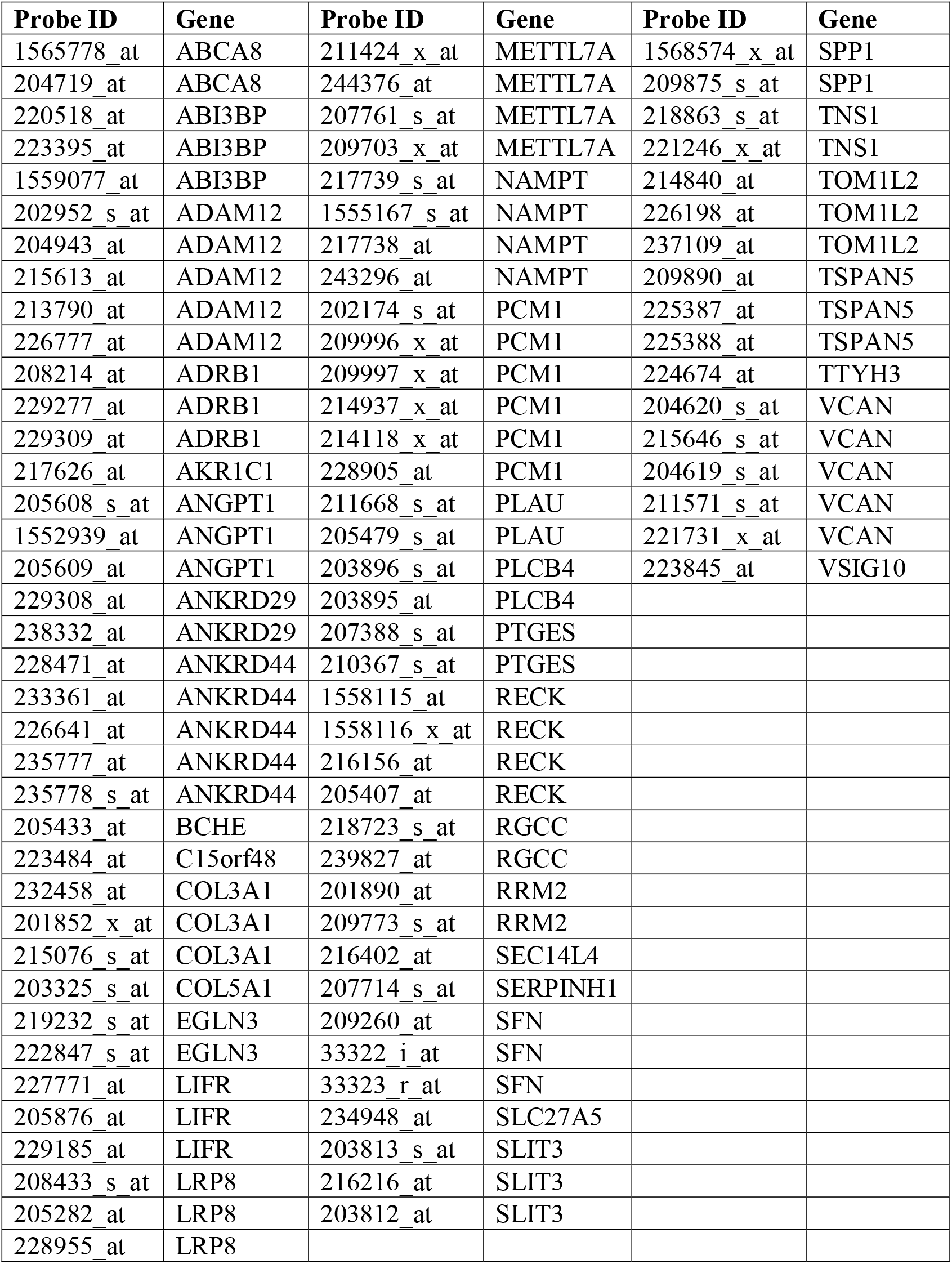
The list of Affymetrix U133 v2.0 probes used in the analysis and the genes to which they map

| Probe ID | Gene | Probe ID | Gene | Probe ID | Gene |
| --- | --- | --- | --- | --- | --- |
| 1565778_at | ABCA8 | 211424_x_at | METTL7A | 1568574_x_at | SPP1 |
| 204719_at | ABCA8 | 244376_at | METTL7A | 209875_s_at | SPP1 |
| 220518_at | ABI3BP | 207761_s_at | METTL7A | 218863_s_at | TNS1 |
| 223395_at | ABI3BP | 209703_x_at | METTL7A | 221246_x_at | TNS1 |
| 1559077_at | ABI3BP | 217739_s_at | NAMPT | 214840_at | TOM1L2 |
| 202952_s_at | ADAM12 | 1555167_s_at | NAMPT | 226198_at | TOM1L2 |
| 204943_at | ADAM12 | 217738_at | NAMPT | 237109_at | TOM1L2 |
| 215613_at | ADAM12 | 243296_at | NAMPT | 209890_at | TSPAN5 |
| 213790_at | ADAM12 | 202174_s_at | PCM1 | 225387_at | TSPAN5 |
| 226777_at | ADAM12 | 209996_x_at | PCM1 | 225388_at | TSPAN5 |
| 208214_at | ADRB1 | 209997_x_at | PCM1 | 224674_at | TTYH3 |
| 229277_at | ADRB1 | 214937_x_at | PCM1 | 204620_s_at | VCAN |
| 229309_at | ADRB1 | 214118_x_at | PCM1 | 215646_s_at | VCAN |
| 217626_at | AKR1C1 | 228905_at | PCM1 | 204619_s_at | VCAN |
| 205608_s_at | ANGPT1 | 211668_s_at | PLAU | 211571_s_at | VCAN |
| 1552939_at | ANGPT1 | 205479_s_at | PLAU | 221731_x_at | VCAN |
| 205609_at | ANGPT1 | 203896_s_at | PLCB4 | 223845_at | VSIG10 |
| 229308_at | ANKRD29 | 203895_at | PLCB4 |  |  |
| 238332_at | ANKRD29 | 207388_s_at | PTGES |  |  |
| 228471_at | ANKRD44 | 210367_s_at | PTGES |  |  |
| 233361_at | ANKRD44 | 1558115_at | RECK |  |  |
| 226641_at | ANKRD44 | 1558116_x_at | RECK |  |  |
| 235777_at | ANKRD44 | 216156_at | RECK |  |  |
| 235778_s_at | ANKRD44 | 205407_at | RECK |  |  |
| 205433_at | BCHE | 218723_s_at | RGCC |  |  |
| 223484_at | C15orf48 | 239827_at | RGCC |  |  |
| 232458_at | COL3A1 | 201890_at | RRM2 |  |  |
| 201852_x_at | COL3A1 | 209773_s_at | RRM2 |  |  |
| 215076_s_at | COL3A1 | 216402_at | SEC14L4 |  |  |
| 203325_s_at | COL5A1 | 207714_s_at | SERPINH1 |  |  |
| 219232_s_at | EGLN3 | 209260_at | SFN |  |  |
| 222847_s_at | EGLN3 | 33322_i_at | SFN |  |  |
| 227771_at | LIFR | 33323_r_at | SFN |  |  |
| 205876_at | LIFR | 234948_at | SLC27A5 |  |  |
| 229185_at | LIFR | 203813_s_at | SLIT3 |  |  |
| 208433_s_at | LRP8 | 216216_at | SLIT3 |  |  |
| 205282_at | LRP8 | 203812_at | SLIT3 |  |  |
| 228955_at | LRP8 |  |  |  |  |

**Supplementary Table 3.**
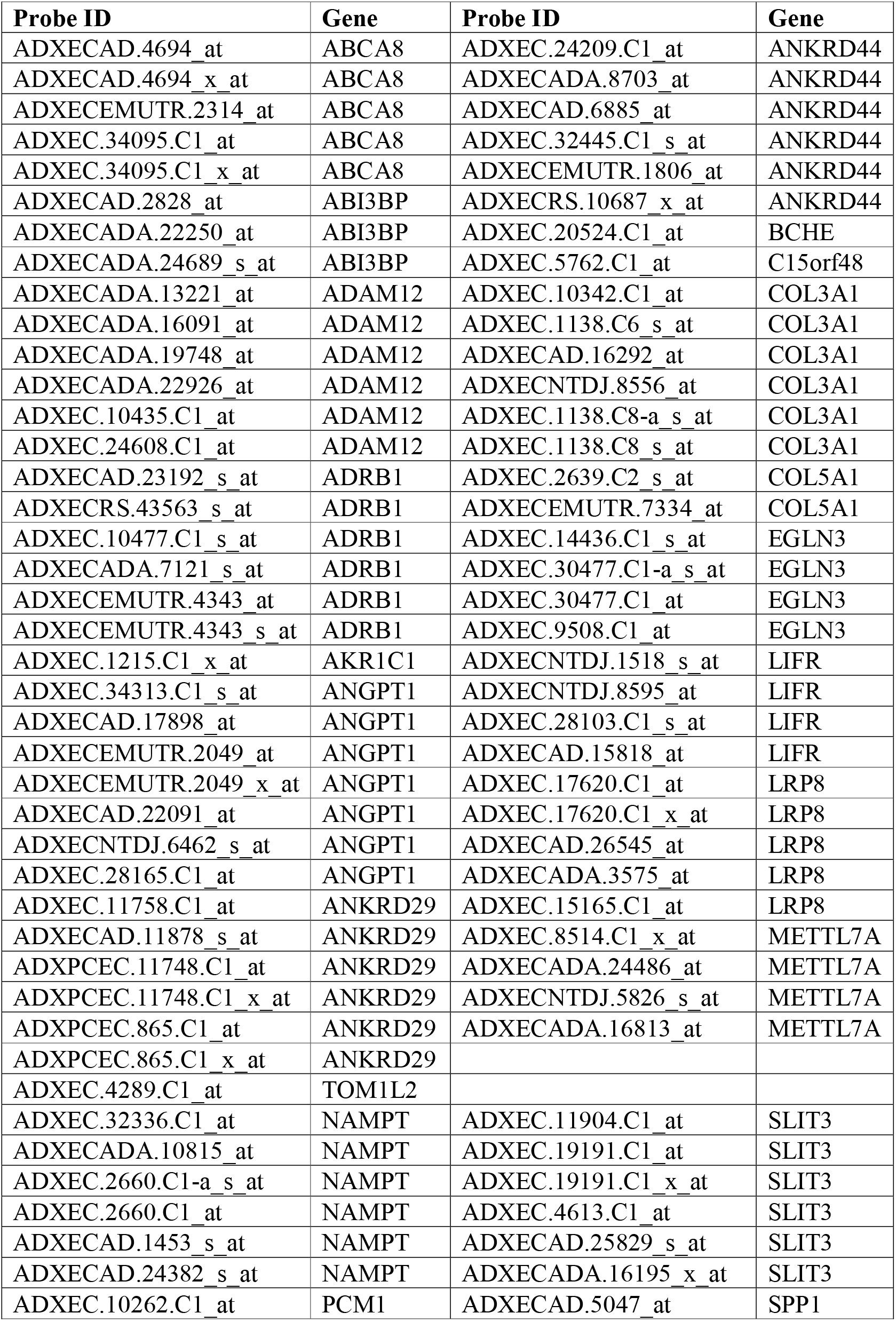
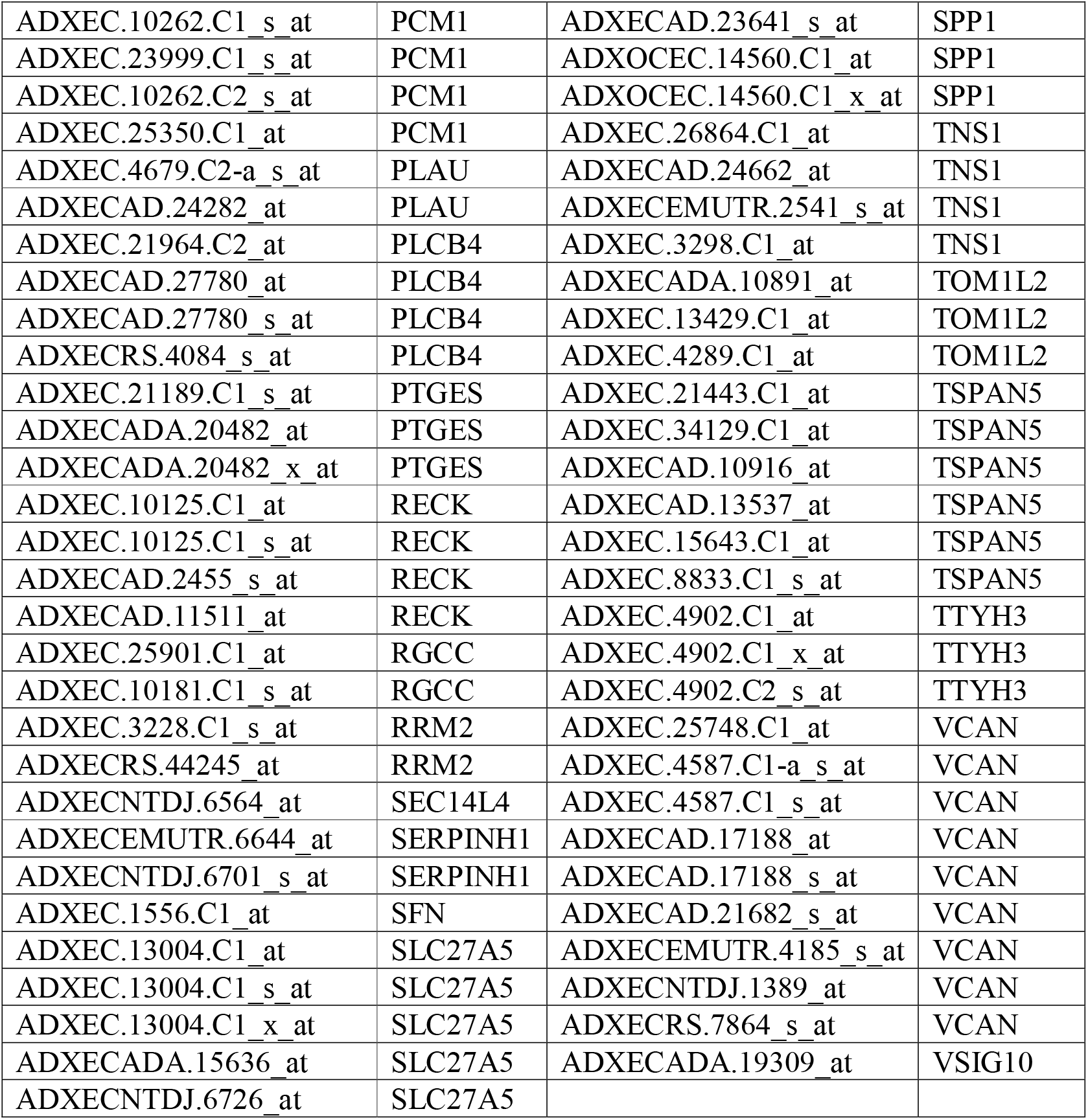
The list of Affymetrix Xcel array probes used in the analysis and the genes to which they map

**Supplementary Table 4.** The list of Agilent Human 4 × 44k v2 array probes used in the analysis and the genes to which they map

| **Probe ID** | **Gene** | **Probe ID** | **Gene** |
| --- | --- | --- | --- |
| A_23_P118615 | ABCA8 | A_33_P3364864 | NAMPT |
| A_33_P3342305 | ABCA8 | A_33_P3364869 | NAMPT |
| A_23_P218858 | ABI3BP | A_33_P3375934 | NAMPT |
| A_33_P3411975 | ABI3BP | A_23_P82950 | PCM1 |
| A_33_P3411980 | ABI3BP | A_24_P555510 | PCM1 |
| A_23_P350512 | ADAM12 | A_23_P24104 | PLAU |
| A_33_P3256848 | ADAM12 | A_33_P3306146 | PLAU |
| A_33_P3310929 | ADAM12 | A_33_P3260634 | PLCB4 |
| A_33_P3310189 | ADRB1 | A_24_P403417 | PTGES |
| A_24_P152968 | AKR1C1 | A_23_P83028 | RECK |
| A_23_P216023 | ANGPT1 | A_33_P3269899 | RECK |
| A_23_P412577 | ANKRD29 | A_24_P225616 | RRM2 |
| A_23_P209347 | ANKRD44 | A_33_P3389286 | SFN |
| A_24_P128713 | ANKRD44 | A_23_P4611 | SLC27A5 |
| A_23_P212050 | BCHE | A_23_P58588 | SLIT3 |
| A_23_P26024 | C15orf48 | A_33_P3391496 | SLIT3 |
| A_24_P935491 | COL3A1 | A_23_P7313 | SPP1 |
| A_23_P83818 | COL5A1 | A_24_P105733 | TNS1 |
| A_33_P3383351 | COL5A1 | A_33_P3209491 | TNS1 |
| A_33_P3256952 | EGLN3 | A_33_P3738458 | TNS1 |
| A_23_P200222 | LRP8 | A_23_P141180 | TOM1L2 |
| A_23_P415021 | METTL7A | A_33_P3325349 | TSPAN5 |
|  |  | A_23_P331895 | TTYH3 |
|  |  | A_23_P144959 | VCAN |

## Footnotes

1 Here, we define the X major PCs as the X most variable PCs that explains 80% of the variation in the data.

2 As the NRF2 metagene expression is a continuous variable, the HR reported in the text of this section is the HR between the upper and lower tertiles of the NRF2 metagene expression.

3 See footnote 2.

